# Rapid differentiation of estrogen receptor status in patient biopsy breast cancer aspirates with an optical nanosensor

**DOI:** 10.1101/2024.03.29.587397

**Authors:** Pooja V. Gaikwad, Nazifa Rahman, Pratyusha Ghosh, Dianna Ng, Ryan M. Williams

## Abstract

Breast cancer is a substantial source of morbidity and mortality worldwide. It is particularly more difficult to treat at later stages, and treatment regimens depend heavily on both staging and the molecular subtype of the tumor. However, both detection and molecular analyses rely on standard imaging and histological method, which are costly, time-consuming, and lack necessary sensitivity/specificity. The estrogen receptor (ER) is, along with the progesterone receptor (PR) and human epidermal growth factor (HER-2), among the primary molecular markers which inform treatment. Patients who are negative for all three markers (triple negative breast cancer, TNBC), have fewer treatment options and a poorer prognosis. Therapeutics for ER+ patients are effective at preventing disease progression, though it is necessary to improve the speed of subtyping and distribution of rapid detection methods. In this work, we designed a near-infrared optical nanosensor using single-walled carbon nanotubes (SWCNT) as the transducer and an anti-ERα antibody as the recognition element. The nanosensor was evaluated for its response to recombinant ERα in buffer and serum prior to evaluation with ER- and ER+ immortal cell lines. We then used a minimal volume of just 10 µL from 26 breast cancer biopsy samples which were aspirated to mimic fine needle aspirates. 20 samples were ER+, while 6 were ER-, representing 13 unique patients. We evaluated the potential of the nanosensor by investigating several SWCNT chiralities through direct incubation or fractionation deployment methods. We found that the nanosensor can differentiate ER-from ER+ patient biopsies through a shift in its center wavelength upon sample addition. This was true regardless of which of the three SWCNT chiralities we observed. Receiver operating characteristic area under the curve analyses determined that the strongest classifier with an AUC of 0.94 was the (7,5) chirality after direct incubation and measurement, and without further processing. We anticipate that further testing and development of this nanosensor may push its utility toward field-deployable, rapid ER subtyping with potential for additional molecular marker profiling.

## Introduction

Breast cancer is the leading cause of cancer-related deaths in women[1], which is often as a result of advanced metastatic disease[2, 3]. Hence, early and accurate diagnosis followed by rapid and appropriate treatment is crucial for decreasing the overall mortality rate. A combination of clinical breast exams, imaging, and biopsy is the standard method for breast cancer detection[4]. Despite advances in high-resolution and non-invasive imaging techniques, early detection is limited by both substantial false negative and false positive rates, cost, and frequency of screening[2, 5]. Therefore, imaging is almost always coupled with lesion biopsy for definitive diagnosis[6]. Fine-needle aspirate biopsy (FNAB) is a widely used diagnostic method due to its minimally invasive nature and cost-effectiveness[7, 8]. Cytology on the aspirate contents allow for diagnosis with high sensitivity and specificity[8]. Immunohistochemistry on FNAB samples is used to determine the molecular phenotype of the lesion, typically evaluating the presence of specific biomarkers of aggressiveness and proliferation[9]. Determinative diagnosis therefore combines information from several analyses and experts, taking a few days in resource-rich healthcare settings and up to several weeks in low-to-middle-income countries[10, 11][12][13]. Given the high cost, time-to-treatment, and issues with diagnostic accuracy, additional methods to improve rapid diagnosis are necessary.

After determining whether the primary lesion is tumor or benign, as well as lymph node or other organ involvement, molecular analyses are used to classify the tumor for treatment and prognostic purposes. This is primarily focused on the receptors for hormones estrogen and progesterone, as well as the epidermal growth factor HER-2[14, 15]. If tumors lack all three of these, they are labeled as triple-negative breast cancer (TNBC), which is substantially more difficult to treat[14, 15]. Molecularly-specific treatment options exist if any of these are present, though the tumor does not respond if treated improperly. Endocrine therapy in estrogen receptor positive (ER+) tumors has proven overall survival and progression-free survival benefit in early stage disease[16, 17] as well tumor reduction in later-stage disease[18-22]. The American Society of Clinical Oncology (ASCO) recommends endocrine therapy as the initial course of treatment for hormone receptor positive metastatic breast cancer[19]. Approximately 15% of breast cancers are triple negative, however, and another 5% are ER-, HER2+[23-26]. Therefore, it is imperative to rapidly determine the presence of these markers.

Single-walled carbon nanotubes exhibit unique characteristics conducive to signal transduction in biological samples. Semiconducting SWCNT are cylindrical fullerenes with near-infrared (NIR) fluorescence emission, which is relatively unabsorbed by biological tissue[27, 28]. There are dozens of semiconducting nanotube chiralities, each with discrete, narrow absorption and fluorescence emission bands and very large Stokes shifts[27], giving rice to the possibility of highly-multiplexed detection via multiple chiral species[29]. Previous work has shown the ability to image nanotubes within biological tissue and in live animals[30, 31]. Thus, loss of signal from biomolecule interference is not a concern with nanotube-based sensing as they are in the so-called ‘tissue window’ of the electromagnetic spectrum[28, 29, 32-34]. Additionally, nanotube fluorescence does not decrease over time as with traditional fluorophores, which has allowed for their frequent and long-term imaging[28, 30, 35].

Nanotube-based optical sensors have previously been developed for diverse classes of molecules and have been adapted for rapid, bedside disease diagnostics. SWCNT fluorescence is dependent on the local environment, thus it can be modulated by binding of antigen following functionalization[36-38]. These analytes include small molecules, nucleic acids, lipids, and proteins, among others[28, 29, 33, 34, 36-39] Thus, SWCNT are ideal transducers of analyte concentrations. Quantitative sensors for biomolecules in live animals have been developed for the majority of these analytes using strategies ranging from direct injection[28, 33] to sensor immobilization in hydrogel matrices or semipermeable membranes[28, 29, 34]. Further, rapid, ex vivo biological sensors using SWCNT have been characterized for multiple disease biomarkers, including lipids, microRNAs, prostate cancer-specific proteins, and ovarian cancer-specific proteins[29, 33, 34, 39, 40]. These have demonstrated functionality in patient serum, urine, cell lines, and ascites.

In this work, we aimed to develop a SWCNT-based rapid detection tool to determine ER status in breast cancer biopsy samples. We took inspiration from prior studies, combining prior detection of cell-surface protein state[40] with molecularly-specific antibody-based detection of cancer biomarkers proteins[34, 39]. We first established a functional SWCNT-based optical sensor for ER in solution, evaluated its response in relevant in vitro models, and then explored the most optimal conditions for detection of ER status in FNAB ex vivo.

### Experimental

#### Preparation of SWCNT-ssDNA

HiPCO (high pressure carbon monoxide)-prepared single-walled carbon nanotubes (SWCNT) (NanoIntegris Technologies, Inc.; Quebec, Canada) were mixed with a single-stranded DNA of the sequence (TAT)_6_-NH_2_, with a 3’primary amine functional group (Integrated DNA Technologies; Iowa, USA) in a 1:2 mass ratio in 0.5 mL 1x phosphate-buffered saline (PBS, Sigma-Aldrich; Missouri, USA). The suspension was sonicated for 1 hour at 40% amplitude while on ice via a 120 W ultrasonicator with 1/8″ probe microtip (Fisher Scientific; New Hampshire, USA). The sonicated suspension was ultracentrifuged for 1 hour at 58,000 x g in 4 ml polycarbonate centrifuge tubes (Beckman Coulter; California, USA) using an Optima Max-XP Ultracentrifuge (Beckman Coulter) fit with an MLA-50 rotor. After ultracentrifugation, the top 75% of the suspension was collected and filtered prior to use with 100 kDa Amicon Ultra 0.5 ml centrifugal filters (Sigma-Aldrich) at 14,000 x g for 15 minutes to remove free ssDNA. After filtration, the solution retained in the filter containing ssDNA-SWCNT was washed two times with 200 μL 1X PBS and centrifugal-filtered again. Finally, the solution containing SWCNT-(TAT)_6_-NH_2_ retained in the filter was suspended in 200 uL PBS.

The concentration of SWCNT-(TAT)_6-_NH_2_was determined by using a V-730 UV–visible absorption Spectrophotometer (Jasco Inc.; Maryland, USA). The concentration of SWCNT was calculated using the value of the absorbance minima near 630 nm (Extinction coefficient = 0.036 L mg^-1^ cm^-1^)[41][42][43].

#### ERα nanosensor synthesis

The amine-modified ssDNA-SWCNT complex was conjugated to a monoclonal ERα antibody (Catalog # 14-9740-82, Invitrogen, Massachusetts, USA) using carbodiimide conjugation chemistry similar to previous studies[41, 42]. Briefly, the carboxylic acid group of the antibodies were activated using 1-ethyl-3-(3-dimethylainopropyl) carbodiimide) (Sigma-Aldrich) and N-hydroxysuccinimide (TCI Chemicals, Oregon, USA), in a 10x and 25x molar excess respectively, for 15 min at 4°C. The reaction was quenched with 1 µL of 2-mercaptoethanol (Sigma-Aldrich). The activated antibodies were mixed with SWCNT-(TAT)_6_-NH_2_in a 1:1 molar ratio of ssDNA to antibody. The reaction mixture was incubated at 4°C on ice for a total of 2 hours, with gentle brief vortexing every 30 minutes. The reaction mixture was dialyzed against deionized water with a 1,000 kDa MWCO filter (Float-A-Lyzer G2; Spectrum Labs, California, USA) at 4°C for 48 hours with two dialysate changes.

#### Physicochemical characterization of the ERα nanosensor

To confirm the successful conjugation of antibody to the ssDNA-nanotube construct, we performed light scattering measurements. Dynamic light scattering (DLS) was performed (Nano-ZS90, Malvern: Worcestershire, United Kingdom) for the nanosensor and SWCNT-(TAT)_6_-NH_2_nanotube complexes as previously described to determine their relative sizes[44, 45].

Electrophoretic light scattering (ELS) (Nano-ZS90, Malvern) was performed to compare the relative ζ-potential of the nanosensor and SWCNT-(TAT)_6_-NH_2_ complex. Data were acquired in triplicate.

#### Characterization of sensor response to ERα in solution

To initially evaluate the response of the ERα nanosensor complex in buffer, we first measured the fluorescence response of 0.5 mg/L nanosensor to 250 nM human ERα (Catalog number NBP2-34478PEP, Novus Biologicals; Colorado, USA) in 1x PBS using a near-infrared (NIR) fluorescent cuvette-based spectrometer (NS MiniTracer; Applied NanoFluorescence, Texas, USA). In triplicate, samples containing 0.5 mg/L nanosensor were spiked with 250 nM ERα. Data was acquired for time points 2, 15, 30, 60, 120, and 180 minutes after addition of ERα antigen. The exposure time used was 3000 ms, with 3 spectra taken and averaged. The instrument excitation wavelength was 638 nm (100 mW laser) and emission was collected from 900-1600 nm via an InGaAs array.

We then measured sensor response in 10% fetal bovine serum (heat-inactivated FBS; Corning, New York, USA). Prior to the addition of antigen, the nanosensor was passivated with 50x mass excess of bovine serum albumin (BSA; Fisher Scientific) compared to the nanosensor for 30 minutes at 4°C as in our prior serum studies[41, 42]. Data was acquired as described above. All experiments were performed in triplicate.

Custom MATLAB code was used to analyze and fit individual nanotube chirality peaks to a pseudo-Voigt model (code available upon request). The analyzed fluorescence emission chiralities were (7,5), (7,6), and (9,5) for excitation with the 638 nm laser source. Fits were ensured to have goodness of fit values (R^2^) greater than 0.98 prior to analyses. Center wavelengths were obtained from each fit, as well as maximum intensity values.

#### Characterization of sensor response in immortalized cell culture models

MDA-MB-231 cells (American Type Culture Collection / ATCC; Virginia, USA) known to lack ERα[46] were cultured in filter-purified 1x DMEM media (Corning) supplemented with 10% FBS and 1X pen/strep (Sigma). HCC1500 (ATCC CRL-2329) cells known to express ERα[47] were cultured in filtered RPMI-1640 media (Corning) with 10% FBS and 1X primocin (InvivoGen; California, USA).

Cells were passaged in T75 culture flasks after reaching 80-90% confluency via 1X TrypLE Express (Fisher) with regular media changes as necessary, with propagation at 37°C with 5% CO_2_ and humidity. Cell counts and viability assays were performed via Trypan blue exclusion and a Countess automated cell counter (Invitrogen).

To evaluate nanosensor response to each cell line, cells were seeded into a 6-well plate with 30,000 cells per well prior to incubation for 3 days. Cells were washed with PBS and fixed with 1 mL 10% formalin (Fisher) for 45 minutes at 37°C. After incubation, cells were scraped and recovered by centrifugation. Cells were resuspended in 500 uL of PBS and diluted with ERα nanosensor solution at a concentration of 0.5 mg/L SWCNT. The final cell count was 0.2 x10^6^ cells/mL and cells were incubated for 15 minutes at 37°C. All samples were centrifuged at 4000xG for 10 minutes, and cell pellets were resuspended in 400 µL PBS.

NIR fluorescence was collected for resuspended triplicates via the MiniTracer cuvette-based spectrometer. Spectral collection parameters were as above for solution-based measurements.

#### Breast cancer patient sample collection

26 total breast cancer samples were collected under Institutional Review Board (IRB) approval at Memorial Sloan Kettering Cancer Center. In total, six breast cancer tumor samples with 0% ERα expression were used (TNBC), with two samples each being derived from three total patients.

For this study, 20 tumor samples were used that were 95% (n=4) or 99% (n=16) ERα positive, two each derived from ten unique patients. Samples were fixed, sectioned, and stained with analysis by an expert pathologist (Ng) to determine whether the tissue was tumor or benign as well as the % breast cancer positivity. Several samples were obtained from different sites of a primary tumor. Biopsy samples were aspirated with a syringe to simulate a fine needle aspirate and suspended in PBS or water. Samples were stored at -80°C and shared with CCNY under Materials Transfer Agreement (MTA).

#### Optimization and characterization of sensor response to patient samples

The ERα nanosensor was passivated with a 50x mass excess of BSA for 30 minutes on ice. The sensor was diluted to 0.5 mg/L at a volume of 90 uL in a 96-well plate (Corning). NIR spectra were acquired using a custom-built high-throughput plate reader spectrophotometer (ClaIR, Photon Etc.; Quebec, Canada). Laser excitation of 655 nm and 730 nm were used sequentially at a power of 1750 mW and integration time 750 ms. Spectra were acquired prior to addition of patient samples, then 10 µL of each breast cancer patient sample were added in triplicated.

Spectra were collected at 2, 30, 60, 90, 120, 150, and 180 minutes post-sample addition. Custom MATLAB code was used to perform background subtraction and fit individual nanotube chirality peaks to a pseudo-Voigt model (code available upon request). The analyzed fluorescence emission chiralities were (7,5), (7,6), and (9,5) for excitation with the 655 nm laser source and (10,2), (9,4), (8,6), (8,7) for excitation with the 730 nm laser source. Chiralities with R^2^ > 0.95 were considered for further analysis. Data are reported as average shift from the pre-sample spectra. As these samples were directly mixed and measured with no further processing, they were designated “unwashed” to differentiate them from further processing below.

We further evaluated whether additional processing was necessary for optimal ERα detection in patient samples. Again, BSA-passivated nanosensor was diluted to 0.5 mg/L in PBS and spectra were obtained in a 96-well plate (ClaIR plate reader). Then, 10 µL of each sample were added in triplicate and incubated for 30 minutes. The samples were transferred to microcentrifuge tubes and centrifuged at 4000xG for 10 minutes. The supernatant was collected, consisting of SWCNT not bound to cells—which we designated as the “washed unbound” fraction. The cell pellet was resuspended in 100 µL PBS, which we designated as the “washed bound” fraction. Each fraction was returned to a 96-well plate and measured with the same parameters as above. Because of the fractionation, samples were dimmer and more difficult to fit. Therefore, only the brightest peaks pertaining to the (7,5), (7,6), and (9,4) chiralities exhibited r^2^ > 0.8 after fitting to the pseudo-Voigt curve and were used for analysis—fits that did not reach that mark were omitted (seven omissions for “washed bound” fraction and six for the “washed unbound” fraction).

#### Statistical analysis and data processing

All statistical analyses were performed in OriginPro 2021 (OriginLab Corporation; Massachusetts, USA). P-values were assigned *** = P < 0.001, ** = P < 0.01, and * = P < 0.05. Comparison of two groups was performed via a two-tailed T-test. Receiver operating characteristic (ROC) curves were plotted in OriginPro to determine the ability of each chirality evaluated and each preparation method to differentiate ER-from ER+ tumor samples. Area under the curve (AUC) was calculated as the primary output of this analysis with asymptotic probability used to determine whether the distinction is significant.

## Results

### Nanosensor synthesis

The ERα nanosensor was constructed using our previously-published carbodiimide chemistry methodology using a monoclonal human anti-ERα antibody[41, 42]. The sensing strategy (**Figure 1A**) was that the fluorescence emission profile of the sensor would change upon recognition of ERα. We first determined that the sensor construct exhibited bright, stable fluorescence upon excitation with 638 nm laser light across the NIR spectrum with peaks corresponding to individual nanotube chiralities characteristic of a bulk SWCNT solution (**Figure 1B**). DLS demonstrated that the correlation coefficient of the nanosensor had a slower decay rate than the unconjugated base ssDNA-SWCNT construct due to a larger hydrodynamic radius caused by the antibody (**Figure 1C**). Additionally, electrophoretic light scattering shows a lesser negative ζ-potential for the ERα nanosensor than that for SWCNT-(TAT)_6_ (**Figure 1D**), further indicating successful sensor synthesis and purification as in prior work[41, 42].

**Figure 1:**
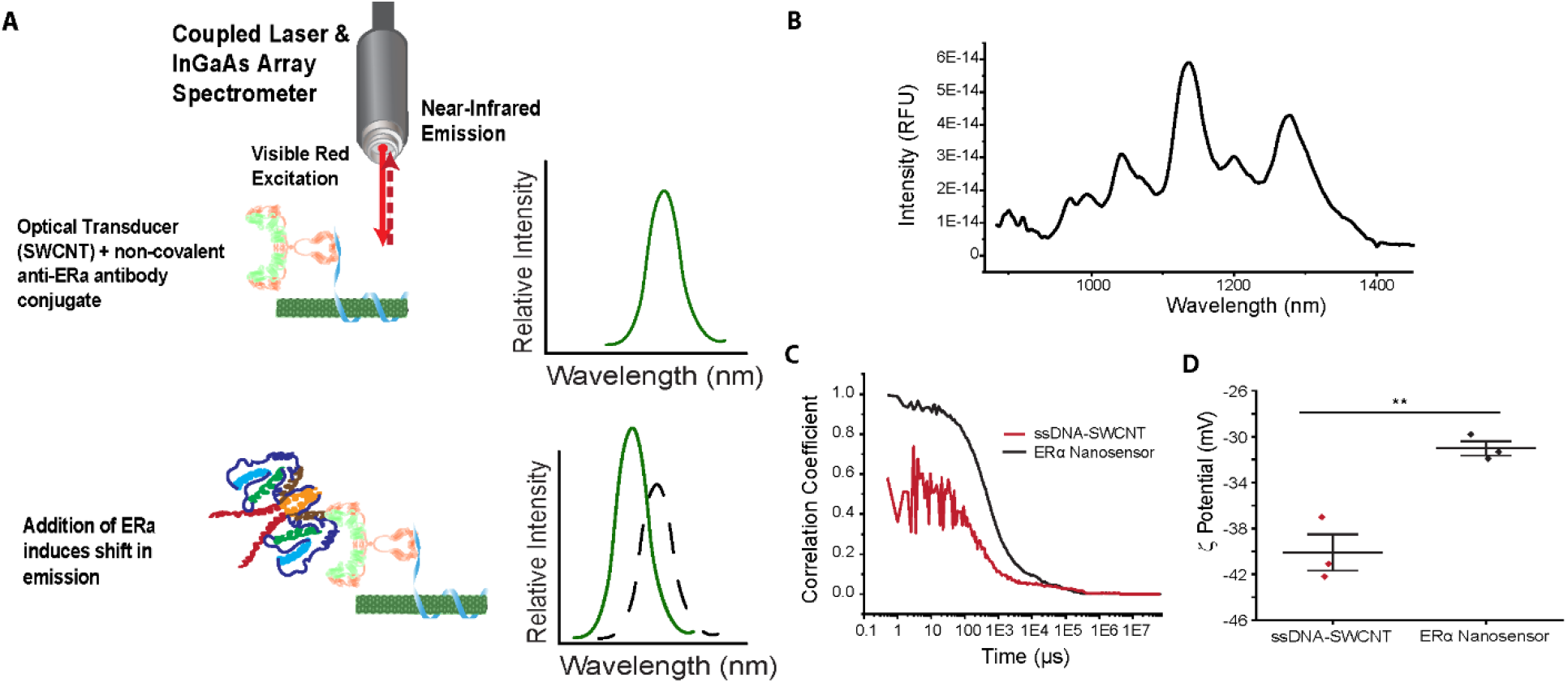
Nanosensor scheme and synthesis. (A) Schematic of the ERα antibody-based NIR fluorescent nanosensor detection concept. (B) NIR fluorescent spectra of the construct nanosensor in PBS. Success of the antibody conjugation to base construct was assessed by (C) comparison of decay in correlation coefficient as a function of time for SWCNT-(TAT)_6_-NH_2_ and ERα antibody (Ab) conjugated to SWCNT- (TAT)_6_, (D) Change in zeta potential for ssDNA-SWCNT compared to the ERα nanosensor. Difference in means = 9.1 mV, p = 0.006.

### Assessing sensor response in buffer

We first evaluated the nanosensor response to 250 nM recombinant ERα in PBS. We found that the center wavelength of the (7,6) chirality of the nanosensor was red-shifted by 3.5 ± 0.2 nm compared to no-protein controls (p = 0.004) (**Figure 2A**). We then evaluated the sensor response to the same recombinant protein in a solution of 10% serum (FBS) to simulate a complex protein-rich environment. The sensor was passivated with BSA as in our prior work to prevent biofouling. We again observed a significant response from the (7,6) chirality of the nanosensor.

**Figure 2:**
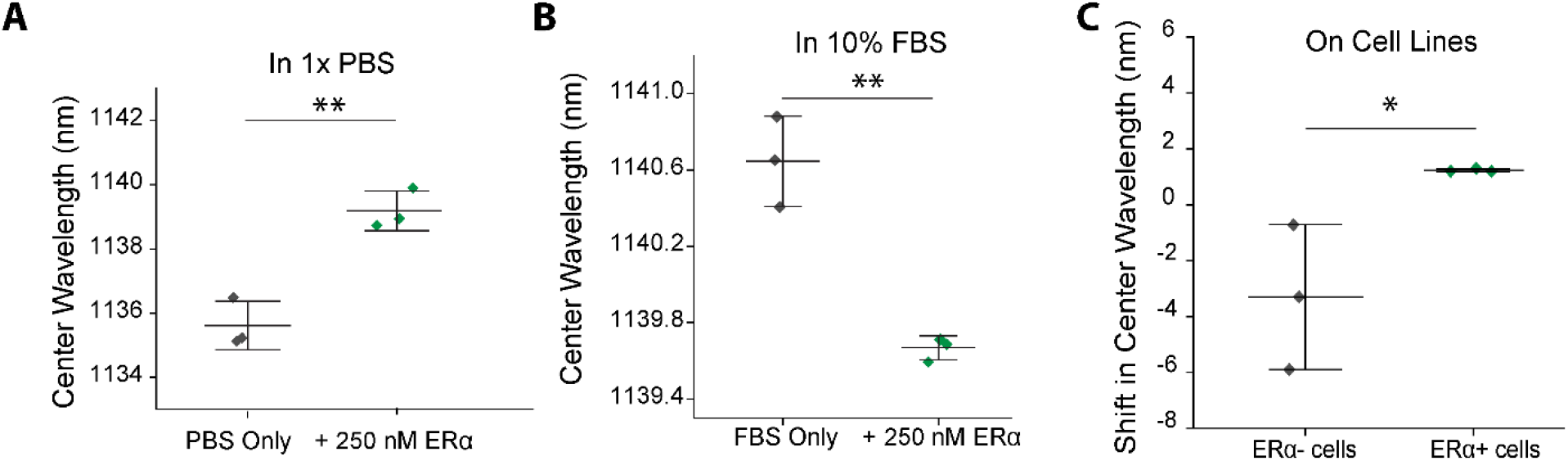
ERα nanosensor response in vitro. (A) Response of the nanosensor (7,6) chirality to 250 nM recombinant ERα in PBS. Difference in means = 3.5 nm, p = 0.004. (B) Response of the nanosensor (7,6) chirality to 250 nM recombinant ERα in 10% FBS. Difference in means = 1 nm, p = 0.002. (C) Change in center wavelength of the (7,5) chirality for the nanosensor incubated with ER- or ER+ cells. Difference in means = 5.5 nm, p = 0.04).

In this case, a blue-shift of 1 ± 0.2 nm was observed (p = 0.01) (**Figure 2B**). It is likely that the center wavelength underwent a blue-shift in 10% FBS as opposed to the red-shift in serum due to rearrangement of passivating BSA or loose protein corona formation.

### Differentiation of ER+ and ER-breast cancer cell lines

We then sought to evaluate the optimal conditions for detection of ERα on the cell surface. We used two cell lines, ERα negative MDA-MB-231 and ERα positive HCC1500 cells. The MDA-MB-231 cell line is a commonly used triple-negative breast cancer (TNBC) cell line, established from a patient with metastatic mammary adenocarcinoma[48]. It lacks expression of estrogen receptor (ER), progesterone receptor (PR), and human epidermal growth factor receptor (HER2) amplification [49][50]. HCC1500 was established from a patient with ductal carcinoma and has estrogen receptor and progesterone receptor expression[51]. Cells were formalin fixed and suspended in PBS to simulate the environment of fixed patient samples.

Cells collected from this method were counted to obtain 12.8 × 10^5^/mL ± 5.6 × 10^5^/mL, whereas a control sample obtained by enzymatic digestion (TrypLE) yielded 15.7 × 10^5^/mL ± 0.93 × 10^5^/mL cells. After 30 minutes of incubation, we observed that the (7,5) chirality demonstrated a 3.3 ± 2.6 nm blue-shift after incubation with ERα -, whereas ERα + cells induced a 1.2 ± 0.02 nm red-shift (p = 0.04) (**Figure 2C**). These results indicate that there is indeed a differential response of the sensor when incubated with TNBC versus ER+ breast cancer cell lines. That the directionality of the difference was a net red-shift of ∼4.5 nm indicates that sensor response was most like that of PBS conditions (**Figure 2A**), as there was no heterogenous formation of protein corona on the sensor.

### Sensor response to breast cancer patient biopsy samples

We then sought to determine the optimal conditions for sensor performance with FNAB-mimic patient samples and to evaluate whether the sensor can differentiate patient tumor samples which are ER-from ER+. To do so, we first measured the sensor fluorescence before and after addition of 10 µL (10% of total solution) of the aspirated patient sample. Each sample was measured in triplicate and the shift in center wavelength at the 90 minute timepoint from prior to addition for each was determined as an average of the three measurements. In total, immunohistochemistry determined that six samples (from three unique patients) were ER-while 20 samples (from ten unique patients) were ER+.

The first condition we evaluated was direct addition with no further processing (**Figure 3A**). We observed three chiralities, the (7,5), (7,6), and (9,4) to determine which were optimal for sample differentiation. We found that for all three chiralities, a red-shift occurred in response to ER-patient samples of 1.8 ± 1.7 nm, 2.3 ± 1.1 nm, and 1.1 ± 0.7 nm, respectively (**Figures 3B-D**). In each case, the average of the ER+ exposed sensor underwent a lower-magnitude red-shift or a blue-shift, with a difference in center wavelength shift of 2.3 nm, 1.6 nm, and 1.2 nm, respectively. Each difference in means was statistically significant (p = 0.0011, 0.0002, and 0.0001, respectively) (**Figures 3B-D**).

**Figure 3:**
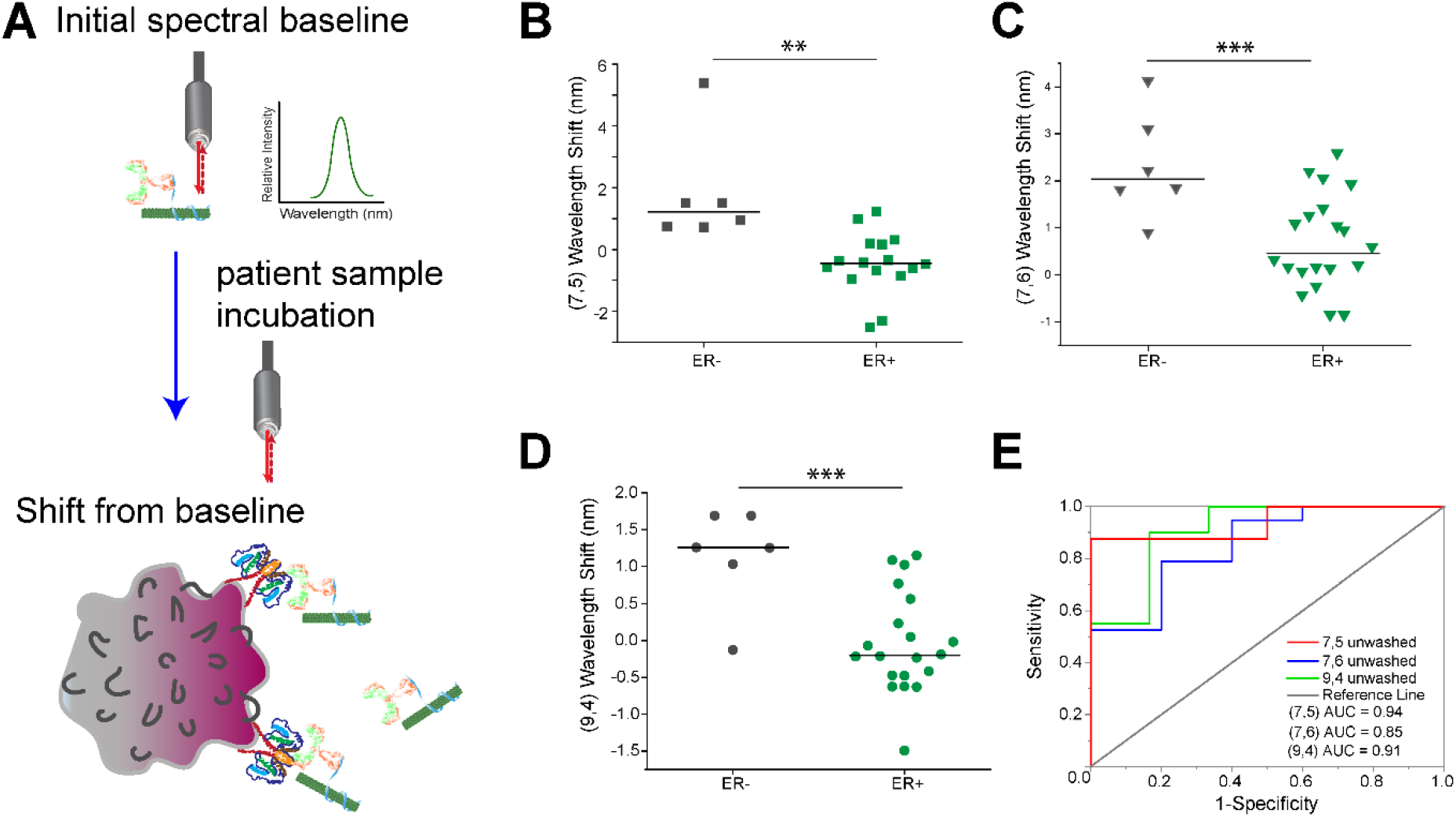
Nanosensor response to aspirated patient biopsy samples. (A) Schematic of direct nanosensor response measurement to ER+ breast cancer patient cells. (B) Change in center wavelength of the nanosensor (7,5) chirality after incubation with ER- or ER+ breast cancer biopsy aspirates. Difference in means = 2.3 nm, p = 0.0011. (C) Change in center wavelength of the nanosensor (7,6) chirality after incubation with ER- or ER+ breast cancer biopsy aspirates. Difference in means = 1.6 nm, p = 0.0002. (D) Change in center wavelength of the nanosensor (9,4) chirality after incubation with ER- or ER+ breast cancer biopsy aspirates. Difference in means = 1.2 nm, p = 0.0001. (E) Receiver operating characteristic evaluation of the ability of each chirality to differentiate ER-from ER+ biopsy samples. AUC is area under the curve. p(7,5) = 0.0020, (7,6) = 0.017, (9,4) = 0.0029.

It is notable that the largest-magnitude shift of the (7,5) chirality gives the most clear distinction between populations. It is particularly exciting that at a cutoff of between 0.4-0.7 nm wavelength shift, only two of the 26 samples would be misclassified (specifically two ER+ samples). From a sensor response point of view, we found it notable that the sensor response more closely resembled the response in 10% FBS, whereas cell line experiments more closely resembled those in PBS. We surmise this is due to the cell line experiments being performed with washed cells in PBS, while patient biopsy samples were aspirates of whole tissue, which included both tumor tissue, extracellular matrix, and other components—more closely resembling the complex protein-rich environment of FBS.

To further evaluate which SWCNT chirality response was the most optimal, we evaluated the area under the curve (AUC) of the receiver operator characteristic (ROC). We found that the AUC for the (7,5), (7,6), and (9,4) responses were 0.94, 0.85, and 0.91, respectively (p = 0.0029, 0.017, and 0.0020, respectively) (**Figure 3E**). We therefore concluded that each of the three evaluated chiralities were able to differentiate between ER- and ER+ tumor tissue with substantial accuracy. However, the (7,5) chirality demonstrated both the largest difference in means of center wavelength shifts as well as the greater AUC.

At the same time, we sought to determine whether samples processed to remove unbound sensor were comparable or optimal to the mixed sample-sensor design above. In this experimental design, sensor that was bound to the cells were separated from sensor remaining in solution via centrifugation after a 30-minute incubation (**Figure 4A**). First, we found that, due to the fractionation protocol, 6-7 of the samples were not sufficiently bright enough for data analysis.

**Figure 4:**
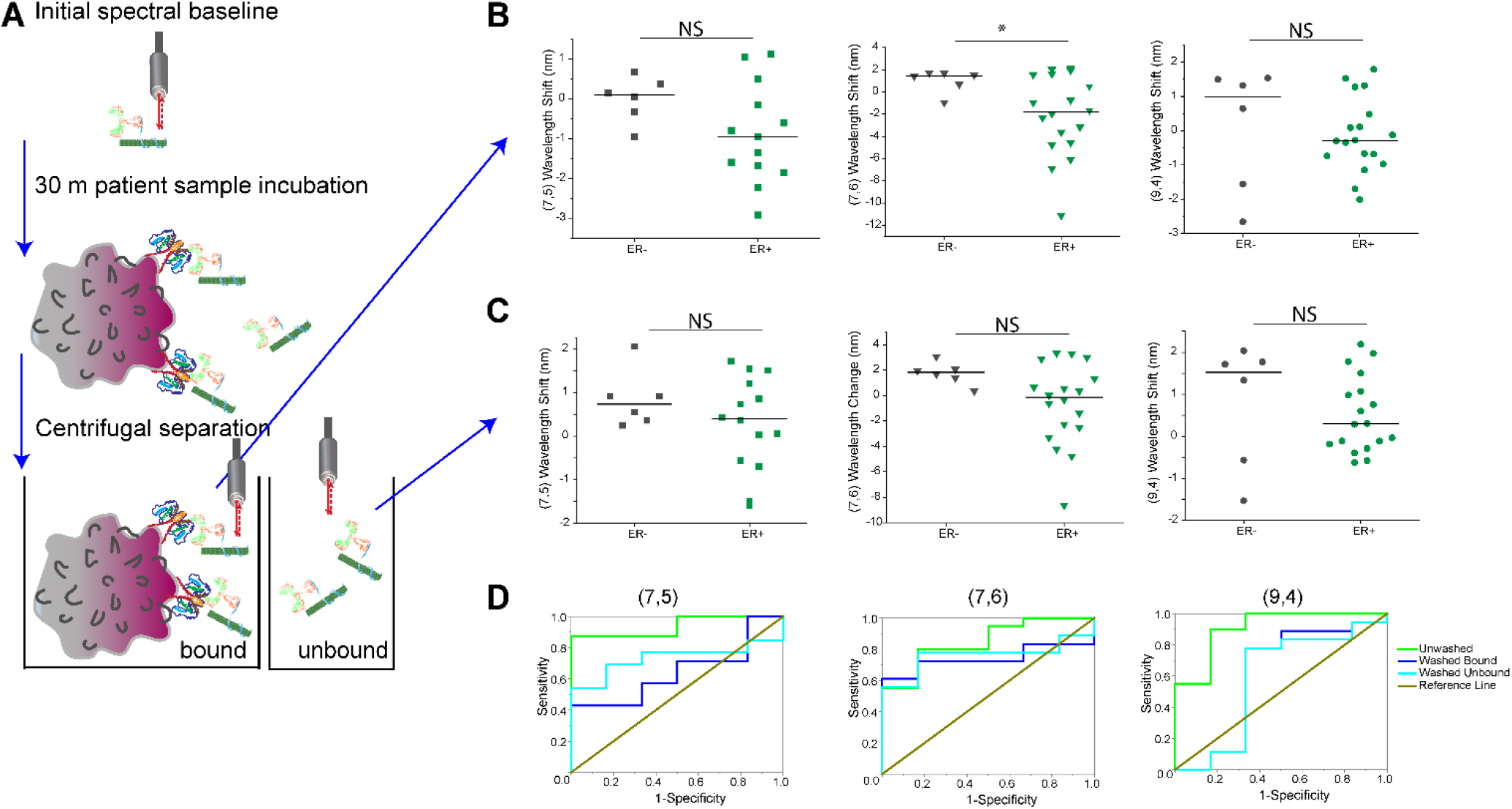
Comparison of fractionation method versus direct measurement. (A) Schematic representing the fractionation method to measure the change in center wavelength for bound vs. unbound nanosensor. (B) Change in center wavelength of the nanosensor each chirality of the “Washed Bound” fraction. Only the (7,6) chirality (center) exhibited a significant change. Difference in the means for this sample = 3.1 nm, p = 0.025. (C) Change in center wavelength of the nanosensor each chirality of the “Washed Unbound” fraction. No differences were statistically significant. (D) ROC comparison of the three measurements (from Figure 3 & 4) and the ability of each chirality to differentiate ER-from ER+ cells. Only the Unwashed method was statistically significant (please refer to Figure 3 for AUC and p values for each).

Of those that were bright enough in the “washed bound” fraction containing sensor bound to cells, we found again found that ER-cells induced a red-shift, while ER+ cells induced a blue-shift. This is consistent with our prior results (**Figure 3**), however only the difference in means of the (7,6) chirality, at 3.1 nm was statistically significant (p = 0.025) (**Figure 4B**). Each of the other chiralities demonstrated a trend in the data showing the same blue-shifted phenomenon for ER+ cells, however several outlier samples existed. This was likely due to low-intensity spectra from having removed much of the sensor.

We then evaluated the sensor change for the “unbound wash” fraction, or the sensor which was not bound to cells. We hypothesized that no difference would be present in these samples as the estrogen receptor is found on the cell surface, though acknowledge that aspiration could sheer some cells, releasing membrane proteins and ER that is not bound to cells. Despite this possibility, we found the expected result of no significant difference in the sensor in solution after incubation with ER- or ER+ FNAB mimics (**Figure 4C**).

Finally, we performed AUC analysis of ROC curves to compare the ability of the sensor to discern ER-from ER+ tumor samples in each of the three preparation conditions: unwashed, washed bound, and washed unbound. For the (7,5) chirality, we found that only the “unwashed” preparation could reliably differentiate the samples (AUC and p values reported above), with neither of the other preparations being better than random chance (p > 0.05) (**Figure 4D**).

Likewise, for the (7,6) and (9,4) chiralities, again only the “unwashed” sample was statistically significant. We therefore concluded that the minimal processing associated with direct measurement and no fractionation was optimal, likely due to improve signal relative to noise.

## Conclusions

In this work, we found that an engineered optical nanosensor for the estrogen receptor can successfully differentiate ER-from ER+ breast cancer biopsies with substantial accuracy. We established optimized methodology aspirate screening with our nanosensor, which operated within 90 minutes and with a minimal sample volume of only 10 µL. We believe this work has substantial potential for rapid differentiation of ER status in the field or in low-resource settings. In our ongoing work, we anticipate additional optimization of the sensor in terms of speed and a more robust change in wavelength to differentiate ER status. We also anticipate multiplexation of this sensor device to evaluate PR and HER2 status at the same time. Finally, we plan to perform larger-scale screens in a broader patient population to approach quantitative stratification of relative abundance of ER positive cells. We are hopeful that this work will reduce the cost and speed necessary for proper diagnosis, allowing proper treatment regimens to be tailored to each patient.

## Acknowledgements

The authors wish to acknowledge all members of the Williams Lab for discussion and feedback. This work was supported by the CCNY-MSKCC Partnership for Cancer Research, Education, and Community Outreach (U54CA132378 / U54CA137788) (RMW and DN) and The City College of New York Grove School of Engineering (RMW). It was also supported by a Dissertation Year Fellowship to P. Gaikwad from the CUNY Graduate Center.

## Notes

### Competing Interest Statement

The authors have declared no competing interest.

